# Macrophage depletion blocks congenital SARM1-dependent neuropathy

**DOI:** 10.1101/2022.02.26.482110

**Authors:** Caitlin B. Dingwall, Amy Strickland, Sabrina W. Yum, Aldrin K. Yim, Jian Zhu, Peter L. Wang, Yurie Yamada, Robert E. Schmidt, Yo Sasaki, A. Joseph Bloom, Aaron DiAntonio, Jeffrey Milbrandt

**Affiliations:** Department of Genetics, Washington University School of Medicine, St. Louis, United States; Children’s Hospital of Philadelphia, Philadelphia, United States; Department of Pathology and Immunology, Washington University School of Medicine, St. Louis, United States; Department of Developmental Biology, Washington University School of Medicine, St. Louis, United States; Needleman Center for Neurometabolism and Axonal Therapeutics, St. Louis, United States; MD-PhD Program, Washington University in St. Louis, St. Louis, United States

**Keywords:** SARM1, axon, neuroinflammation, macrophages, NAD^+^, SARMopathy

## Abstract

Axon loss contributes to many common neurodegenerative disorders. In healthy axons, the axon survival factor NMNAT2 inhibits SARM1, the central executioner of programmed axon degeneration. We identified two rare *NMNAT2* missense variants in two brothers afflicted with a progressive neuropathy syndrome. The polymorphisms result in amino acid substitutions, V98M and R232Q, which reduce NMNAT2 NAD^+^-synthetase activity. We generated a mouse model of the human syndrome and found that *Nmnat2*^V98M^/*Nmnat2*^R232Q^ compound-heterozygous CRISPR mice survive to adulthood but develop progressive motor dysfunction, peripheral axon loss, and macrophage infiltration. These disease phenotypes are all SARM1-dependent. Remarkably, macrophage depletion therapy blocks and reverses neuropathic phenotypes in *Nmnat2*^V98M/R232Q^ mice, identifying a SARM1-dependent neuroimmune mechanism as a key driver of disease pathogenesis. These findings demonstrate that SARM1 induces an inflammatory neuropathy and highlight the potential of immune therapy to treat this rare syndrome and other neurodegenerative conditions associated with NMNAT2 loss and SARM1 activation.

## INTRODUCTION

Axon loss is one of the earliest pathological hallmarks and likely the initiating event in many common neurodegenerative disorders^1–3^. Programmed axon degeneration is an active, genetically encoded subcellular self-destruction pathway executed by the pro-degenerative NADase enzyme SARM1 (sterile alpha and Toll/interleukin-1 receptor motif-containing)^4,5^. In a healthy axon, SARM1 activity is restrained by the NAD^+^ biosynthetic enzyme NMNAT2, which converts NMN and ATP into NAD^+6^. NMNAT2 is a highly labile protein produced in the soma and trafficked into the axon^7^. Nerve injury blocks axonal transport and leads to rapid depletion of axonal NMNAT2^8^, causing NMN buildup and NAD^+^ loss. Recent breakthroughs led to the discovery that SARM1 is activated by an increase in the NMN to NAD^+^ ratio^9^. NMN and NAD^+^ can both bind at an allosteric site in the enzyme’s N-terminus to differentially regulate its conformation, and hence, the activation state of SARM1^9–11^. The ratio of NMN/NAD^+^ rises after NMNAT2 loss, favoring SARM1 activation and subsequent axon degeneration^9^. Genetic deletion of NMNAT2 alone is perinatal lethal; however, when combined with SARM1 deletion, mice are viable and resistant to injury-induced axon degeneration, suggesting that an essential role of NMNAT2 is to inhibit SARM1^12^.

SARM1 is the central executioner of a cell-autonomous axon degeneration program. Upon activation, SARM1, an NAD^+^ hydrolase, depletes axonal NAD^+^ levels, culminating in metabolic crisis and axon fragmentation. Acute injury is the best-understood trigger of pathological axon degeneration, inducing distal axon loss^5,13–18^. Loss of SARM1 is protective in models of chemotherapy-induced peripheral neuropathy (CIPN)^14,16^ and traumatic brain injury (TBI)^15^. However, programmed axon degeneration is also common in chronic neurodegenerative disease models that do not include acute axonal injury^14,16,19–25^, suggesting a role for subacute SARM1 activation in the pathogenesis of a wide range of chronic neurodegenerative conditions.

Evidence for the involvement of the NMNAT2/SARM1 axon degeneration pathway in chronic disease has emerged from studying rare human patient mutations. Indeed, in a model of Leber congenital amaurosis type 9 (LCA9), SARM1 depletion rescues photoreceptor cell death caused by loss of the nuclear NMNAT isoform, *NMNAT1*^*26*^. Furthermore, *NMNAT2* mutations were identified in a still born fetus with fetal akinesia deformation sequence and two sisters suffering from a mild polyneuropathy^27,28^. The first direct evidence of SARM1 mutations in human disease emerged with the discovery of rare hypermorphic *SARM1* alleles in a subset of ALS patients^29,30^. However, until the present study, the field lacked a mechanistically defined model of a SARM1-dependent, chronic axonopathy (termed “SARMopathy”).

Here we examine two brothers that presented in early childhood with recurring Guillain– Barré-like episodes requiring mechanical ventilation combined with severe, progressive peripheral neurodegeneration. Whole exome sequencing revealed they are both compound heterozygotes for two rare missense variants in the *NMNAT2* gene, each inherited from one of their parents. We created a mouse that harbors both mutant *NMNAT2* alleles and found that this model recapitulates key features of the human syndrome. NMNAT2 is the endogenous inhibitor of SARM1 in axons; thus, defects in NMNAT2 can trigger aberrant activation of SARM1. Indeed, we find that these disease phenotypes are all SARM1-dependent. While SARM1 is best understood as the executioner of a cell-autonomous axo-destructive program, here we make the surprising discovery that SARM1 induces axon degeneration via induction of non-cell-autonomous macrophage activation.

Macrophages play complex roles as both pro-degenerative and pro-restorative immune cells. Indeed, neuroinflammation has been referred to as a “double-edged sword” as it can have both beneficial and deleterious effects on the nervous system^32^. On one hand, in the peripheral nervous system, macrophages play a necessary role in facilitating axon regeneration through the clearance of myelin and axonal debris after nerve injury^33–35^. Indeed, macrophage depletion hampers axon regeneration after peripheral nerve injury^36–39^. However, macrophages and their CNS counterparts, microglia, are also drivers of disease in several common neurodegenerative diseases, including multiple sclerosis (MS)^40^, chemotherapy-induced neuropathy (CIPN)^41^, and Alzheimer’s Disease (AD)^42^. In these disease models, depletion of macrophages and microglia can mitigate disease phenotypes, suggesting a conserved pro-degenerative role for phagocytes in human neurological disease.

Collectively, our data establish *NMNAT2* variants as the genetic basis of a human neuropathy and demonstrate an unexpected role for SARM1 as a driver of neuroinflammation in the peripheral nervous system. In this model, we find that macrophage depletion early in the disease course can block the development of neuropathy, and remarkably, treatment after symptom onset can reverse neuropathy phenotypes. Our study provides the first mouse model of a chronic, *injury-independent* SARM1-dependent axonopathy (“SARMopathy”) with which to test axon degeneration-blocking treatments. This will be of substantial benefit and high clinical relevance as the field uncovers an ever-growing list of chronic neurodegenerative diseases that involve SARM1 activation. Importantly, our work uncovers a SARM1-dependent non-cell-autonomous mechanism of axon loss and highlights macrophage depletion as a potent axo-protective therapeutic strategy.

## RESULTS

### Rare missense variants in *NMNAT2* cause hereditary neuropathy

Patients 1 and 2 are brothers from non-consanguineous, healthy parents of African American ancestry. No further members of the extended family are known to be affected. Patient 1 was born following an uneventful pregnancy. Development was normal and the patient acquired the ability to walk before the onset of illness. At age 13 months, he experienced an acute episode of hypotonia, weakness, and respiratory failure requiring hospitalization and mechanical ventilation. Electrophysiology testing (nerve conduction studies and electromyography) at the time of symptom onset showed features of multifocal, sensory and motor neuropathy thought to be consistent with Guillain-Barre Syndrome. After treatment with intravenous immune globulin (IVIG) and steroids, he regained some motor function and was taken off ventilatory support but exhibited residual weakness.

In subsequent years he developed a unique sensorimotor syndrome comprised of *both* chronic and episodic features. Episodic attacks are frequently contemporaneous with infection and include severe neuropathic pain, worsening erythromelalgia, flaccid quadriparesis, and respiratory failure requiring mechanical ventilation. During these episodes, electrophysiological testing shows a complete absence of sensory and motor responses. In between episodes, the patient experiences a chronic, progressive motor-predominant peripheral neuropathy. Currently 25-years-old, Patient 1 is cognitively normal and attends college. Electrophysiology testing indicates a predominantly motor axonal neuropathy. He is wheelchair-dependent, exhibits progressive scoliosis, poor weight gain, and has severe combined proximal and distal muscle atrophy with predominantly distal muscle weakness (**Fig. 1a**). Muscle ultrasound revealed fatty, fibrotic tissue replacement of muscle, consistent with chronic neuropathy^43^. The patient also experiences recurring neuropathic pain, erythromelalgia, bilateral optic atrophy, and tongue fasciculation. Cranial and spinal MRI are normal except for mildly prominent extra-axial spaces. At age 15, a head CT showed mild diffuse parenchymal atrophy or pseudoatrophy. Also at 15, the patient had a normal echocardiogram and EKG testing indicated a possible enlarged atrium. Pulmonary function was tested, and spirometry demonstrated normal lung function.

**Figure 1:**
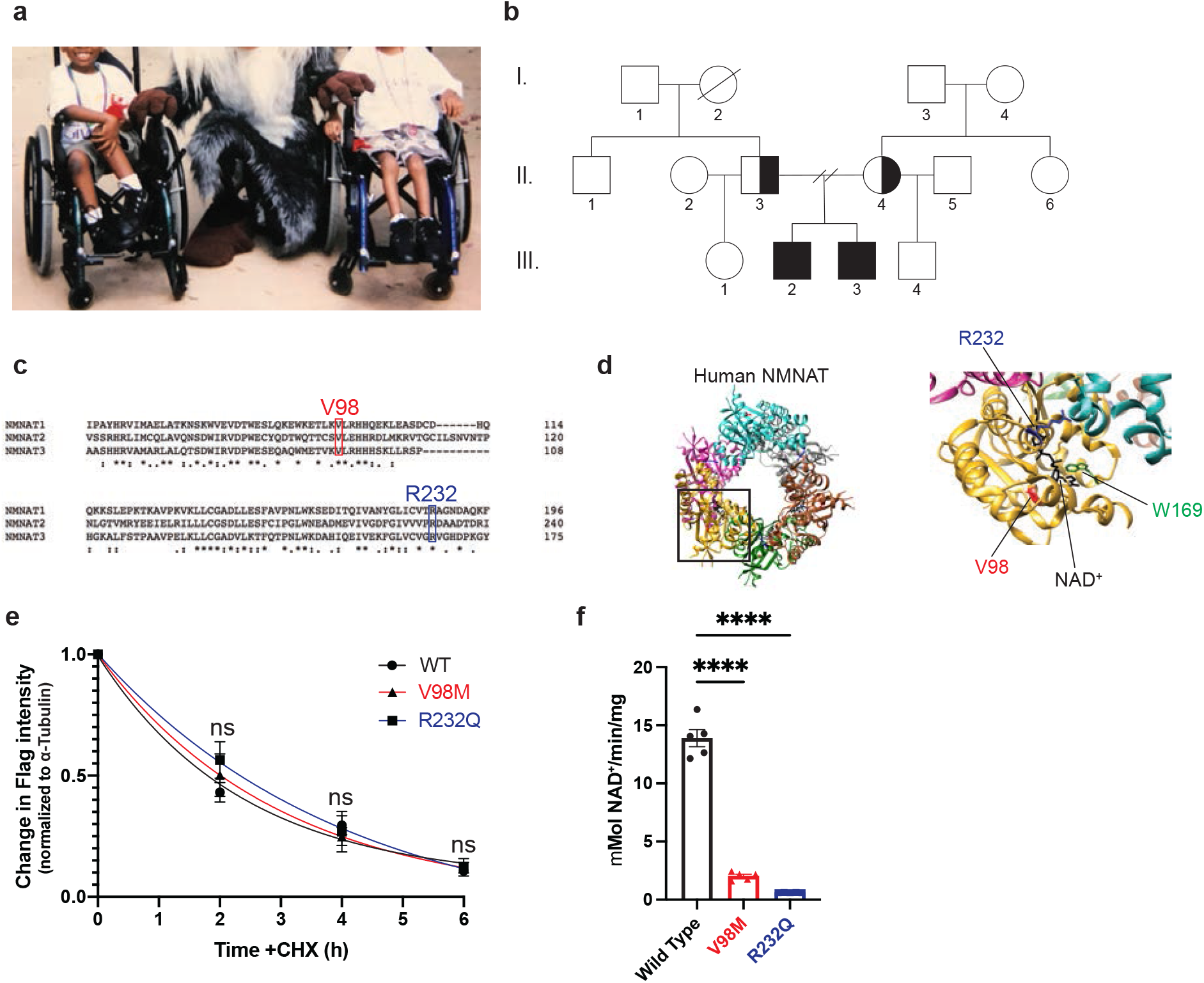
Identification of compound heterozygous *NMNAT2* variants in two brothers with relapsing-remitting neuropathy. **a**, Clinical features of two brothers suffering a unique sensorimotor neuropathy syndrome. **b**, WES showed that both brothers carry two extremely rare missense mutations in the *NMNAT2* gene (c.292G>A. c.695G>A), each inherited from one of their parents. **c**, Both residues are conserved in all three human *NMNAT* isoforms. **d**, Schematic of NMNAT structure. The location of the patients’ missense variants is noted in red (V98) and blue (R232). W169 (green) is the catalytic residue. **e**, Relative turnover rates for Flag-NMNAT2 (wild-type, V98M, R232Q) after cycloheximide (CHX) addition. Changes in Flag-NMNAT intensity were normalized to α-tubulin (loading control) and calculated as a proportion of the intensity of the 0h untreated band for each variant. One-phase decay curves were fitted to the data using non-linear regression. **f**, NMNAT activity assay was performed on purified NMNAT2 at 37°C. NAD^+^ production at steady state (10 min) was used to calculate the NMNAT activity. All data are presented as mean +/- SEM from n=5 independent experiments. Statistical significance was determined by two-way ANOVA with multiple comparisons. ns: not significant, ****p<0.0001.

Patient 2 was born 3 years after Patient 1 following an uneventful pregnancy. Patient 2 met early developmental milestones but experienced symptoms similar to his sibling prior to learning to walk. His first episode of severe weakness requiring mechanical ventilation occurred at 11 months. Patient 2’s clinical course has been virtually identical to his brother’s with very similar symptoms and degree of impairment.

Whole-exome sequencing was performed on the brothers and their parents to identify candidate variants that may cause the patients’ disease. Both affected patients share rare, compound heterozygous variants [c. 695G>A (p.Arg232Gln) and c.292G>A (p.Val98Met)] in *NMNAT2*. Each parent was found to be heterozygous for one of the two variants identified in the patients (**Fig. 1b**). R232Q was previously identified as a loss-of-function variant associated with Fetal Akinesia Deformation Sequence and occurs in a region of NMNAT2 involved in substrate binding^28^. V98M is a novel NMNAT2 variant of unknown significance. Both variants occur at residues that are conserved in all three human NMNAT isoforms (**Fig. 1c-d**).

### V98M reduces NMNAT2 NAD^+^ synthetase activity but not protein stability

We sought to elucidate the functional consequences of the NMNAT2 variant alleles. To investigate whether NMNAT2^V98M^ altered protein stability, we used immunoblotting to compare its relative half-life to NMNAT2^R232Q^ and NMNAT2 in transfected HEK cells. In line with prior studies of NMNAT2 half-life^8^, protein synthesis blockade leads to a rapid drop in NMNAT2 protein levels. Turnover rates of NMNAT2^V98M^ and NMNAT2^R232Q^ are not significantly different from that of control NMNAT2 (**Fig. 1e**).

NMNAT2 is an NAD^+^ synthesizing enzyme, and this activity is required for its function as an axon survival factor. To investigate whether NMNAT2^V98M^ has impaired enzymatic activity, we purified recombinant Strep-tagged NMNAT2 proteins using affinity chromatography^44^. NMNAT2^V98M^ has 14.6% of the NAD^+^ synthesis activity of wild-type NMNAT2 at 37°C, whereas NMNAT2^R232Q^ is 4.4% as active as the wild-type enzyme, in agreement with previous findings^28^ (**Fig. 1f**). Collectively, these data demonstrate that these *NMNAT2* variants disrupt enzymatic function, which may underly their pathogenicity.

### *Nmnat2*^*V98M/R232Q*^ mice develop progressive motor neuropathy

To study the pathological effects of the V98M and R232Q *NMNAT2* mutations found in these patients, we used CRISPR-induced mutagenesis to create mice with mutations *Nmnat2*^*V98M*^ or *Nmnat2*^*R232Q*^ (see methods). Mice heterozygous for these individual mutations were mated to generate mice with compound heterozygous mutations *Nmnat2*^*V98M*^ and *Nmnat2*^*R232Q*^ (henceforth referred to as *Nmnat2*^V98M/R232Q^ mice). These mutant mice were viable with no evidence of embryonic or perinatal lethality. As patients with compound heterozygous variants in *NMNAT2* exhibit a chronic, motor-predominant peripheral neuropathy, we searched for similar phenotypes in the *Nmnat2*^*V98M/R232Q*^ mice. Starting at two months, we assayed muscle strength using an inverted screen test and found that the mice exhibit age-dependent, progressive muscle weakness (**Fig. 2a**). The human disorder involves predominantly distal muscle atrophy; therefore, we assayed hindlimb grip strength in the mutant mice. We observed a decline in distal muscle strength (**Fig. 2b**). Gait defects manifest in *Nmnat2*^*V98M/R232Q*^ mice as young as six months old, concomitant with progressive lower limb muscular atrophy. The majority of mice display severe hindlimb wasting and difficulty walking by 9-12 months of age (**Supp. Video 1**). Notably, while the human patients have episodic neuropathic pain, we did not elicit a nociceptive defect in tail flick testing of *Nmnat2*^*V98M/R232Q*^ mice (**Fig. 2c**).

**Figure 2:**
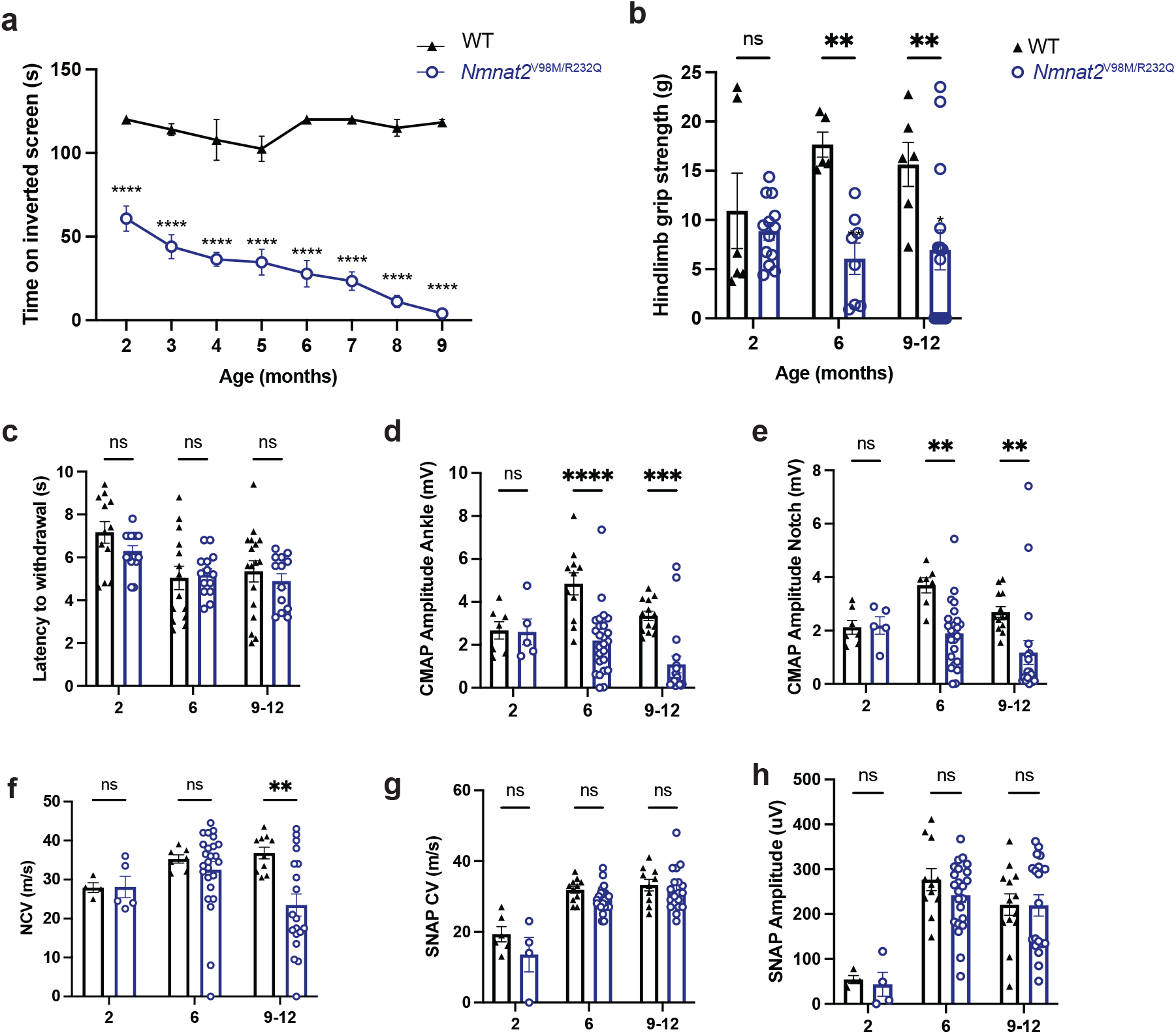
*Nmnat2*^*V98M/R232Q*^ mice have behavioral and electrophysiologic features consistent with a motor neuropathy. **a**, Average time suspended from an inverted screen (max. 120s) for WT or *Nmnat2*^V98M/R232Q^ mice. **b**, Hindlimb grip strength for WT or *Nmnat2*^*V98M/R232Q*^ mice at 2, 6, and 9-12 months of age. **c**, The average time it takes for WT or *Nmnat2*^*V98M/R232Q*^ mice to remove their tails from a 55°C hot water bath at 2, 6, and 9-12 months of age. **d-e**, CMAP amplitude of WT and *Nmnat2*^*V98M/R232Q*^ mice at the ankle (d) and sciatic notch (e) at 2, 6, and 9-12 months of age. **f**, NCV (m/s) of sciatic nerves of WT or *Nmnat2*^*V98M/R232Q*^ mice at 2, 6, and 9-12 months of age. **g**, SNAP CV (m/s) of sciatic nerves of WT or *Nmnat2*^*V98M/R232Q*^ mice at 2, 6, and 9-12 months of age. **h**, SNAP amplitude (μV) of sciatic nerves of WT and *Nmnat2*^*V98M/R232Q*^ mice at 2, 6, and 9-12 months of age. All data are presented as mean +/- SEM. Statistical significance was determined by two-way ANOVA with multiple comparisons. ns: not significant, *p<0.05, **p<0.01, ***p<0.001, ****p<0.0001.

### *Nmnat2*^*V98M/R232Q*^ mice have electrophysiologic features consistent with a motor neuropathy

The decreased muscle strength observed in *Nmnat2*^*V98M/R232Q*^ mice suggests motor neuron dysfunction. We measured motor fiber function using compound muscle action potential (CMAP) amplitudes and found significant deficits in *Nmnat2*^*V98M/R232Q*^ mice. The abnormalities worsen with age, suggesting that motor axon numbers progressively diminish in parallel with decreasing overall strength (**Fig 2d-e**). Next, we tested motor nerve conduction velocity (NCV). A decrease in NCV early in disease without altered CMAP amplitudes is indicative of demyelination, whereas a progressive drop in NCV concomitant with low CMAP amplitudes indicates large-diameter axon loss. The NCV in young *Nmnat2*^*V98M/R232Q*^ mice is normal, indicating that the disease is primarily an axonal neuropathy; however, NCV does decrease with age, likely due to the eventual loss of large diameter axons (**Fig. 2f**). Electrophysiologic sensory testing demonstrated that large, myelinated sensory axons are not affected in *Nmnat2*^*V98M/R232Q*^ mice (**Fig 2g-h**). Pain is transmitted by small and thinly myelinated fibers; thus nerve conduction studies are typically unaffected^45^. Rather, intraepidermal nerve fiber density (IENFD) analysis is a more sensitive measure of small fiber loss. In agreement with normal nociceptive function, we found that IENFD is unaffected in *Nmnat2*^*V98M/R232Q*^ mice (**Fig. S1**). Altogether, these data demonstrate that *Nmnat2*^*V98M/R232Q*^ mice have a motor axonal neuropathy, consistent with the chronic electrophysiological features of human patients with *NMNAT2*-associated neuropathy.

### *Nmnat2* variants cause progressive axon loss and muscle wasting in mice

To further characterize the disease process in *Nmnat2*^*V98M/R232Q*^ mice, we used light microscopic analysis to examine the pathology of select peripheral nerves including the sciatic (a mixed nerve), femoral (primarily motor), and sural (primarily sensory) in 2-month, 6-month, and 9-12-month-old mice. The sciatic and femoral nerves exhibit severe, progressive axon loss. In contrast, we did not observe progressive axon loss in the sural nerve; however, total axon area was modestly different from wild-type control mice at 9-12 months of age (**Fig. 3a-c**). Myelin thickness is not affected in any of the nerves we examined (**Fig. S2**).

**Figure 3:**
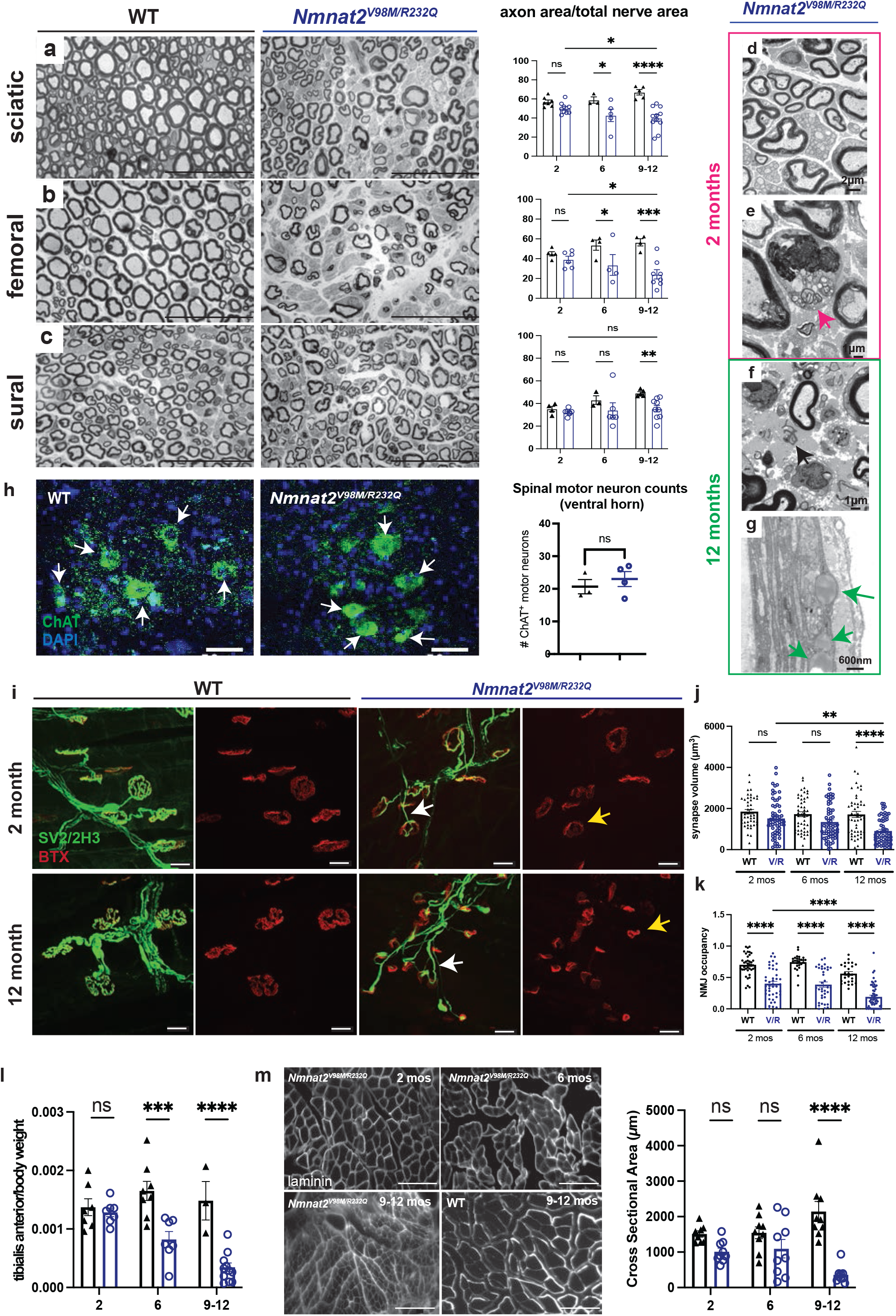
*Nmnat2* variants cause progressive axon loss and muscle wasting in mice. **a-c**, Representative images of sciatic (a), femoral (b), and sural (c) nerves in 9-12-month-old *Nmnat2*^*V98M/R232Q*^ or WT mice. Percent axon area as a ratio of total nerve area is calculated to the right. Scale bar represents 50μm. **d**, The appearance of the sciatic nerve at two months shows a dense population of large and small myelinated axons with little intervening extracellular space. **e**, Macrophage containing axonal and myelin debris are found in the endoneurial space of two-month-old *Nmnat2*^*V98M/R232Q*^ sciatic nerves. **f**, The sciatic nerve at 12 months shows patches of marked axon loss with increased collagen and wispy processes of Schwann cells. Scattered macrophages with axonal and myelin debris are identified. **g**, Presence of large perineurial droplets of neutral fat. **h**, Representative images of ChAT immunostaining in 12-month-old *Nmnat2*^*V98M/R232Q*^ or WT mouse spinal cord (ventral horn), scale bar represents 50μm. Quantification of number of ChAT^+^ motor neuron cell bodies in the ventral horn to the right. **i**, Representative images of 2- and 12-month-old mouse NMJs stained for synaptic vesicle 2/neurofilament (green) and bungarotoxin (red). Scale bar represents 20μm. **j**, Quantification of synapse volume for WT and *Nmnat2*^*V98M/R232Q*^ NMJs. **k**, Quantification of overlap between presynaptic area vs. postsynaptic area for WT and *Nmnat2*^*V98M/R232Q*^ NMJs at 2, 6, and 12 months of age. **l**, Average tibialis anterior weight/body weight for WT and *Nmnat2*^*V98M/R232Q*^ mice in 2, 6, and 9-12-month-old mice. **m**, Representative images of laminin immunofluorescence in *Nmnat2*^*V98M/R232Q*^ mouse tibialis anterior muscles at 2, 6, and 9-12 months of age. WT mouse tibialis anterior muscle at 12 months of age shown for comparison. The apparent fuzziness shown in the representative image of *Nmnat2*^*V98M/R232Q*^ 9–12-month muscle was a consistent genotype-dependent finding reflecting diffuse laminin staining. Quantification of muscle fiber cross-sectional area to the right. Scale bar represents 150μm. All data are presented as mean +/- SEM. Statistical significance was determined by Student’s unpaired t-test or two-way ANOVA with multiple comparisons. ns: not significant, *p<0.05, ***p<0.001, ****p<0.0001.

We next performed electron microscopic analysis on 2-month and 12-month-old *Nmnat2*^*V98M/R232Q*^ sciatic nerves. The appearance of the sciatic nerve at two months is normal and shows a dense population of large and small myelinated axons with little intervening extracellular space (**Fig. 3d**). Schwann cells and macrophages containing axonal and myelin debris are found in the endoneurial space (**Fig. 3e, arrow**). The sciatic nerve at 12 months shows patches of marked axon loss with increased collagen and wispy processes of Schwann cells (**Fig. 3f, arrow**). Following axon degeneration, perineurial cells take up lipid droplets from myelin breakdown^46^; indeed, we find large perineurial droplets of neutral fat in the 12-month sciatic nerve (**Fig. 3g, arrow**). To confirm that the observed peripheral defects are not due to motor neuron cell death, we immunostained spinal cords of 12-month-old wild-type and *Nmnat2*^*V98M/R232Q*^ mice for the motor neuron marker choline acetyltransferase (ChAT) (**Fig. 3h**). Motor neuron cell numbers in the ventral horn are equivalent between genotypes, and thus, axon loss in *Nmnat2*^*V98M/R232Q*^ nerves is likely not due to motor neuron cell death. Taken together, these pathological features demonstrate a progressive peripheral axonal neuropathy.

We next examined the neuromuscular junctions in the hindpaw lumbrical muscles of *Nmnat2*^*V98M/R232Q*^ mice. We find the NMJ endplate size is slightly diminished in mutant mice even as early as 2 months of age (**Fig. 3i, yellow arrows**). The NMJ postsynaptic volume continues to progressively diminish over time, consistent with loss of presynaptic inputs (**Fig. 3i-j**). Apposition of the presynaptic nerve terminal and the postsynaptic endplate is a major determinant of NMJ functionality. Indeed, the ratio of overlap between presynaptic vesicles and the underlying acetylcholine receptor clusters (NMJ occupancy) is reduced in NMJs of *Nmnat2*^*V98M/R232Q*^ 2-month-old mice and decreases over time (**Fig. 3i, k**). In addition, endplate complexity is decreased in 12-month-old *Nmnat2*^*V98M/R232Q*^ mice, whereas the prototypical pretzel-like endplate structure is still observed in 2-month-old *Nmnat2*^*V98M/R232Q*^ mice (**Fig. 3i, yellow arrows**). Alterations in endplate size and complexity suggest repeated episodes of denervation and reinnervation. At steady state, the majority of *Nmnat2*^*V98M/R232Q*^ endplates appear partially innervated (**Fig. 3i**), however, almost all pre-terminal motor axons are abnormally thin and smooth (**Fig. 3i, white arrows**), a hallmark of sprouting axons^47^. Sprouting is frequently observed in NMJs of motor neuron disease models and is evidence of continual axonal degeneration and regeneration^47–50^. Taken together, these data indicate that decreased NMNAT2 activity causes progressive degeneration of terminal axons at the NMJ, consistent with a chronic motor neuropathy.

The patients with compound heterozygous variants in *NMNAT2* have both proximal and distal weakness with predominantly distal muscle atrophy rendering them wheelchair-bound. Loss of nerve terminals at the NMJ results in muscle fiber denervation and eventual muscle atrophy. *Nmnat2*^*V98M/R232Q*^ mice exhibit a progressive reduction in hindlimb muscle mass (**Fig. 3l**) that correlates with decreased fiber cross-sectional area in the tibialis anterior muscle (**Fig. 3m**). Together, these results confirm that the decreased activity of the *NMNAT2* mutants leads to axon loss and subsequent muscle denervation and atrophy. Importantly, the *Nmnat2*^*V98M/R232Q*^ mouse model recapitulates chronic motor features of the human syndrome, providing strong evidence that the *NMNAT2* variants are indeed pathogenic.

### Neuronal SARM1 is required for *Nmnat2*^*V98M/R232Q*^ neuropathy

SARM1 is a pro-degenerative enzyme activated by binding the NAD^+^ precursor NMN at its allosteric site^9^. NMNAT2 converts NMN to NAD^+^ thereby preventing the buildup of NMN and its interaction with SARM1. In *Nmnat2* KO mice, the increase in NMN leads to axon projection abnormalities and perinatal death; however, *Nmnat2/Sarm1* double KO mice are viable and completely resistant to injury-induced programmed axon degeneration^12^. To determine whether SARM1 is activated in *Nmnat2*^*V98M/R232Q*^ mice, we first monitored nerve levels of cADPR, a specific biomarker of SARM1 NAD^+^ hydrolase activity^31^. Metabolites were isolated from the sciatic nerve of two-month-old *Nmnat2*^*V98M/R232Q*^ mice and analyzed by LC/MS-MS. We find that cADPR levels are elevated 8-fold compared to wild-type and this increase is fully SARM1-dependent (**Fig. 4a**), demonstrating that SARM1 is activated even at this early stage of the disease and suggesting that cADPR is likely an excellent biomarker for syndromes involving chronic SARM1 activation.

**Figure 4:**
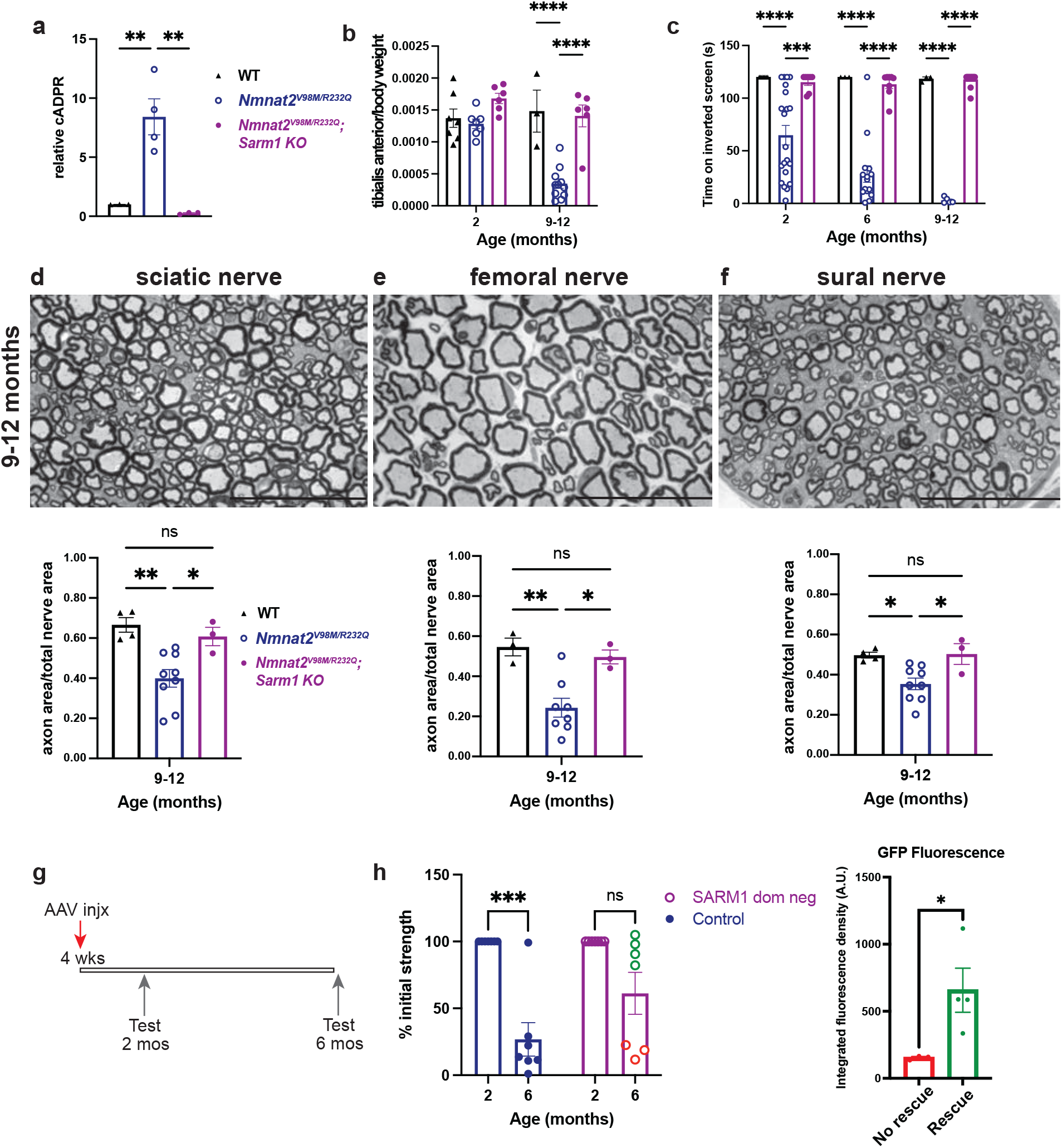
*Sarm1* knockout rescues neuropathy caused by *Nmnat2* variants in mice. **a**, Relative cADPR levels in sciatic nerves of 2-month-old WT, *Nmnat2*^*V98M/R232Q*^, and *Nmnat2*^*V98M/R232Q*^; *Sarm1 KO* mice. Values normalized to WT cADPR levels (set to 1). Statistical significance determined by a Student’s unpaired t-test, **p<0.01. **b**, Average tibialis anterior weight by body weight for WT, *Nmnat2*^*V98M/R232Q*^, and *Nmnat2*^*V98M/R232Q*^; *Sarm1 KO* mice in 2 and 9-12-month-old mice. **c**, Average time suspended from an inverted screen (max. 120s) for WT, *Nmnat2*^*V98M/R232Q*^, and *Nmnat2*^*V98M/R232Q*^; *Sarm1 KO* mice. **d-f**, Representative images of sciatic (d), femoral (e), and sural (f) nerves in 9-12-month-old *Nmnat2*^*V98M/R232Q*^; *Sarm1 KO* mice. Scale bar represents 50μm. Percent axonal area as a ratio of total nerve area is calculated below each corresponding nerve. All data are presented as mean +/- SEM. Statistical significance was determined by two-way ANOVA with multiple comparisons. ns: not significant, *p<0.05, **p<0.01, ***p<0.001, ****p<0.0001. **g**, Schematic of AAV-SARM1-DN gene therapy experiment. **h**, Percent initial performance on inverted screen test at 2 months and 6 months of age for EGFP (control) or SARM1 dominant-negative injected *Nmnat2*^*V98M/R232Q*^ mice. Statistical significance within treatment group was determined by a Student’s paired t-test, ns: not significant, **p<0.01. Statistical significance between treatment groups was determined by a Student’s unpaired t-test, ns: not significant, *p<0.05.

Next, we mated the *Nmnat2*^*V98M/R232Q*^ mutant to *Sarm1* KO mice to generate *Nmnat2*^*V98M/R232Q*^; *Sarm1* KO mice. In contrast with *Nmnat2*^*V98M/R232Q*^ (*Sarm1* WT) mice, *Nmnat2*^*V98M/R232Q*^; *Sarm1* KO mice do not develop motor function deficits (**Fig. 4b-c**). We also performed morphological analysis of the sural, femoral, and sciatic nerves of *Nmnat2*^*V98M/R232Q*^ and *Nmnat2*^*V98M/R232Q*^; *Sarm1* KO mice. As with the functional studies, loss of *Sarm1* prevents axon degeneration even in the oldest *Nmnat2*^*V98M/R232Q*^ mice (**Fig. 3a, 4d-i**). These results confirm that the phenotypes associated with these pathogenic *NMNAT2* variants are SARM1-dependent and do not arise secondary to neomorphic functions of the mutant NMNAT2 enzymes. These results suggest that *Nmnat2*^*V98M/R232Q*^ mice will be useful for testing treatment strategies for progressive neurodegenerative disease involving chronic SARM1 activation.

The development of small molecule and gene therapy SARM1 inhibitors is underway^22,51^; indeed, we previously showed that adeno-associated virus (AAV)-mediated, neuron-specific expression of a potent SARM1 dominant-negative (SARM1-DN) blocks pathologic axon degeneration in models of acute nerve injury^22,52^. With the discovery that the *Nmnat2*^*V98M/R232Q*^ motor neuropathy is SARM1-dependent, we next tested whether this SARM1 gene therapy approach could similarly block disease in these mice. One-month-old *Nmnat2*^*V98M/R232Q*^ mice received intrathecal injections of AAV virions (6 × 10^11^) expressing SARM1-DN-EGFP or EGFP alone (control) under control of a synapsin promoter (**Fig. 4j**). We assayed inverted screen performance at 2 months and 6 months for both groups and determined therapeutic efficacy by comparing 6-month to 2-month performance for each mouse. *Nmnat2*^*V98M/R232Q*^ mice injected with EGFP (control) display a ∼73% decline in strength by 6 months of age (p<0.0001) (**Fig. 4k**). In contrast, *Nmnat2*^*V98M/R232Q*^ mice injected with SARM1-DN exhibit a 39% decline (n.s., p>0.05) at 6 months of age (**Fig. 4k**). Importantly, SARM1-DN rescue of strength defects is dependent on extent of viral infection and transgene expression in the spinal cord. For example, higher expression correlates with higher endpoint performance, whereas lower expression correlates with lower endpoint performance (**Fig. 4k, Fig. S3**). These results demonstrate that neuron-specific high-level expression of SARM1-DN potently protects *Nmnat2*^*V98M/R232Q*^ mice from developing motor deficits, demonstrating that neuron-autonomous SARM1 activity is pathogenic in *Nmnat2*^*V98M/R232Q*^ mice.

### Macrophages orchestrate *Nmnat2*^*V98M/R232Q*^ neuropathy

After acute nerve injury, macrophages infiltrate the lesion and phagocytose axonal and myelin debris, clearing the injury site and promoting axonal regeneration^33–35^. In models of chronic neurodegenerative disease, macrophages and their CNS counterpart, microglia, play complex immunomodulatory roles as both pro- and anti-inflammatory mediators in disease^41,42,53–59^. We immunostained central (spinal cord) and peripheral (sciatic nerve) nervous tissue with antibodies to the activated macrophage marker CD68^60^. There is a notable absence of CD68^+^ cells in central and peripheral tissues from wild-type mice (**Fig. 5a, c**). In contrast, in sciatic nerves of 2-month-old *Nmnat2*^*V98M/R232Q*^ mice, CD68^+^ activated macrophages (**Fig. 5e**) are present, concomitant with a trend toward elevated CD64^+^CD11b^+^ total nerve macrophages (**Fig. 5f**). Significantly fewer CD68^+^ macrophages are observed in *Nmnat2*^*V98M/R232Q*^ mice lacking SARM1 at the same age (**Fig. 5d**), and the number of total nerve macrophages (CD64^+^CD11b^+^) in these animals was similar to wild-type (**Fig. 5f**). CD68^+^ macrophages are not detected in the spinal cords of *Nmnat2*^*V98M/R232Q*^ mice (**Fig. 5b**), consistent with the predominantly peripheral nervous system defects in the patients. NAD^+^ biosynthesis plays a role in programming macrophage immune responses^61^; thus, we examined whether intrinsic defects exist in *Nmnat2*^*V98M/R232Q*^ macrophages due to impaired NMNAT2 activity. However, we found no differences in either basal or antigen-induced activation between wild-type and *Nmnat2*^*V98M/R232Q*^ -derived murine peritoneal macrophages (**Fig. S3**). This is consistent with findings in other disorders indicating that signals within the neural microenvironment shape macrophage phenotype activation^62–66^ as demonstrated by increased numbers of activated macrophages in *Nmnat2*^*V98M/R232Q*^ sciatic nerves.

**Figure 5:**
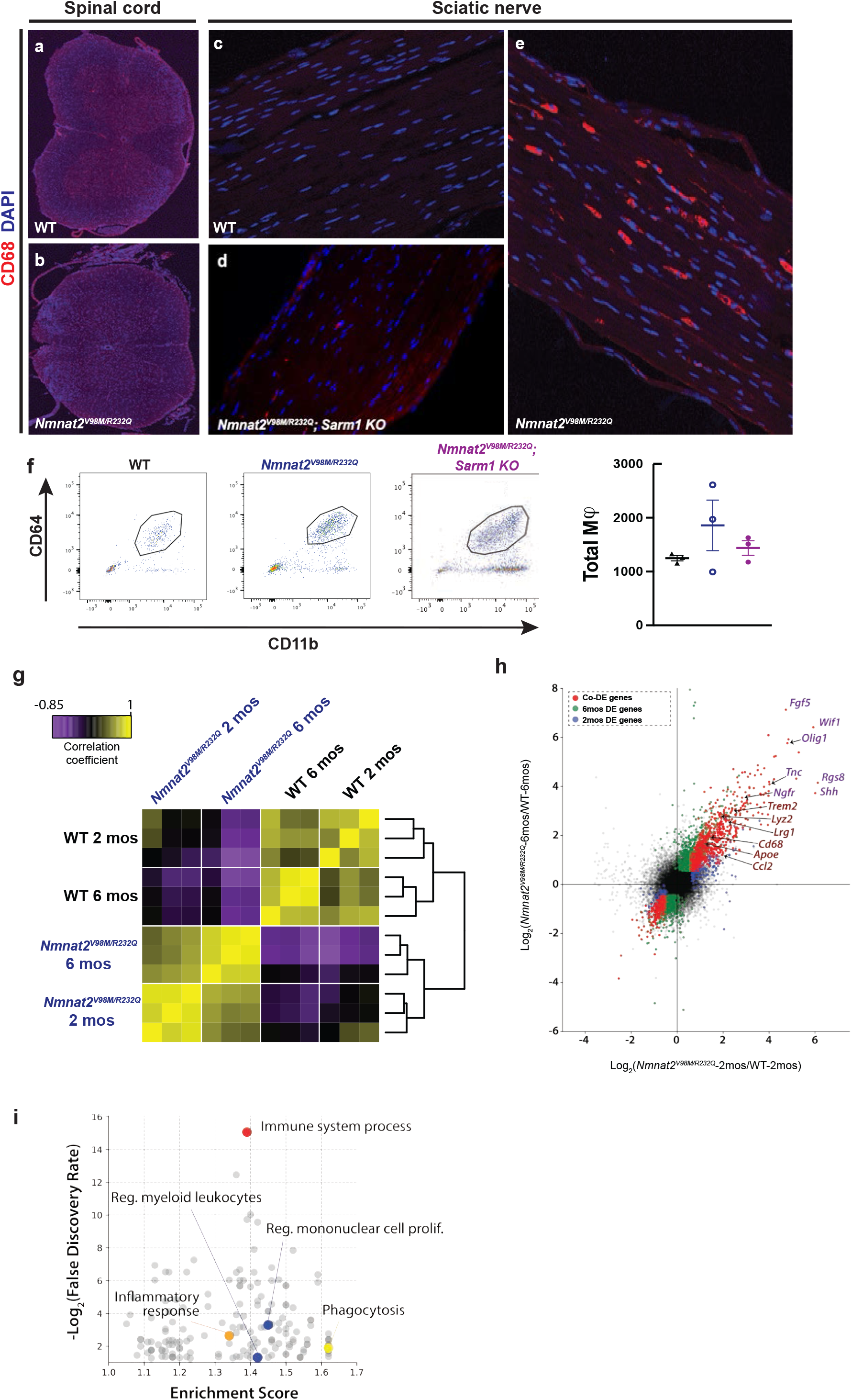
Activated macrophages accumulate in the peripheral nervous system of *Nmnat2*^*V98M/R232Q*^ mice. **a-e**, Representative images of CD68 immunofluorescence and DAPI signal in the spinal cord (a-b) and sciatic nerves (c-e) of 2-month-old WT, *Nmnat2*^*V98M/R232Q*^, and *Nmnat2*^*V98M/R232Q*^; *Sarm1 KO* mice. **f**, Representative scatter plots and quantification of fluorescence-activated cell sorting of total sciatic nerve macrophages (CD64^+^ CD11b^+^) in 2-month-old WT, *Nmnat2*^*V98M/R232Q*^, and *Nmnat2*^*V98M/R232Q*^; *Sarm1 KO* mice. **g**, Sample correlation plot showing global transcriptomic analysis and hierarchical clustering of sciatic nerve macrophages from WT and *Nmnat2*^*V98M/R232Q*^ mice. Each box represents one replicate. **h**, Volcano plot of significant co-differentially expressed genes (DEGs) in 2-month and 6-month *Nmnat2*^*V98M/R232Q*^ sciatic nerves highlighting activated macrophage markers and repair Schwann cell signatures. **i**, GO analysis of genes enriched in 6-month *Nmnat2*^*V98M/R232Q*^ sciatic nerves.

We performed bulk RNA Sequencing (RNA-Seq) on *Nmnat2*^*V98M/R232Q*^ and wild-type (control) mouse sciatic nerves at early and late disease stages to capture dynamic changes in the glial and immune cell milieu (**Fig. 5g**). Global transcriptomic analysis revealed similarities within *Nmnat2*^*V98M/R232Q*^ nerves at both 2 months and 6 months, clustering more closely to each other than wild-type nerves. Closer inspection revealed sets of activated macrophage signatures^62^, including *Cd68, Trem2, Apoe, Lrg1* and *Ccl2*, up-regulated in both 2-month and 6-month old *Nmnat2*^*V98M/R232Q*^ mouse sciatic nerves (**Fig. 5h**), consistent with our observations of CD68^+^ nerve-associated macrophages in *Nmnat2*^*V98M/R232Q*^ mice. Repair Schwann cell (SCs) markers^67–69^ such as *Fgf5, Shh, Ngfr*, and *Olig1* are also upregulated, suggesting a dedifferentiation program of myelinating SCs. Gene ontology (GO) analysis showed significant enrichment of genes related to the immune response, inflammation, and phagocytosis signatures in both 2-month and 6-month-old *Nmnat2*^*V98M/R232Q*^ mice (**Fig. 5i & Supp. Tables 1-2**). Altogether, these data demonstrate chronic activation of peripheral nervous system macrophages and an ongoing Schwann cell repair program in this mouse model.

Increased numbers of activated macrophages in the nerves of the *Nmnat2*^*V98M/R232Q*^ mice raised the question of whether they are playing a beneficial or destructive role in the disorder. We used a macrophage depletion strategy using colony stimulating factor 1 receptor (CSF1R) blockade^71,72^ to evaluate the role of macrophages in *Nmnat2*^*V98M/R232Q*^ mice. The mice were treated with CSF1R monoclonal antibody (or, isotype control antibody: IgG) every three weeks beginning at one month of age, and assessed using motor function tests at 2, 3, and 4 months of age (**Fig. 6a)**. The efficacy of the macrophage depletion treatment was confirmed by immunostaining for CD68 in sciatic nerves (**Fig. S4**). Remarkably, macrophage depletion completely blocks the development of muscle strength defects for the duration of the experiment. In contrast, *Nmnat2*^*V98M/R232Q*^ mice treated with isotype control antibody (IgG) continue to exhibit poor motor function (**Fig. 6b**). Morphological examination of the femoral (predominantly motor) nerve shows that macrophage depletion significantly rescues axon loss in the femoral nerve (**Fig. 6c**), consistent with its ability to prevent motor function deficits. Together, these data show that macrophages promote axon degeneration that leads to motor function deficits in this mouse neuropathy model.

**Figure 6:**
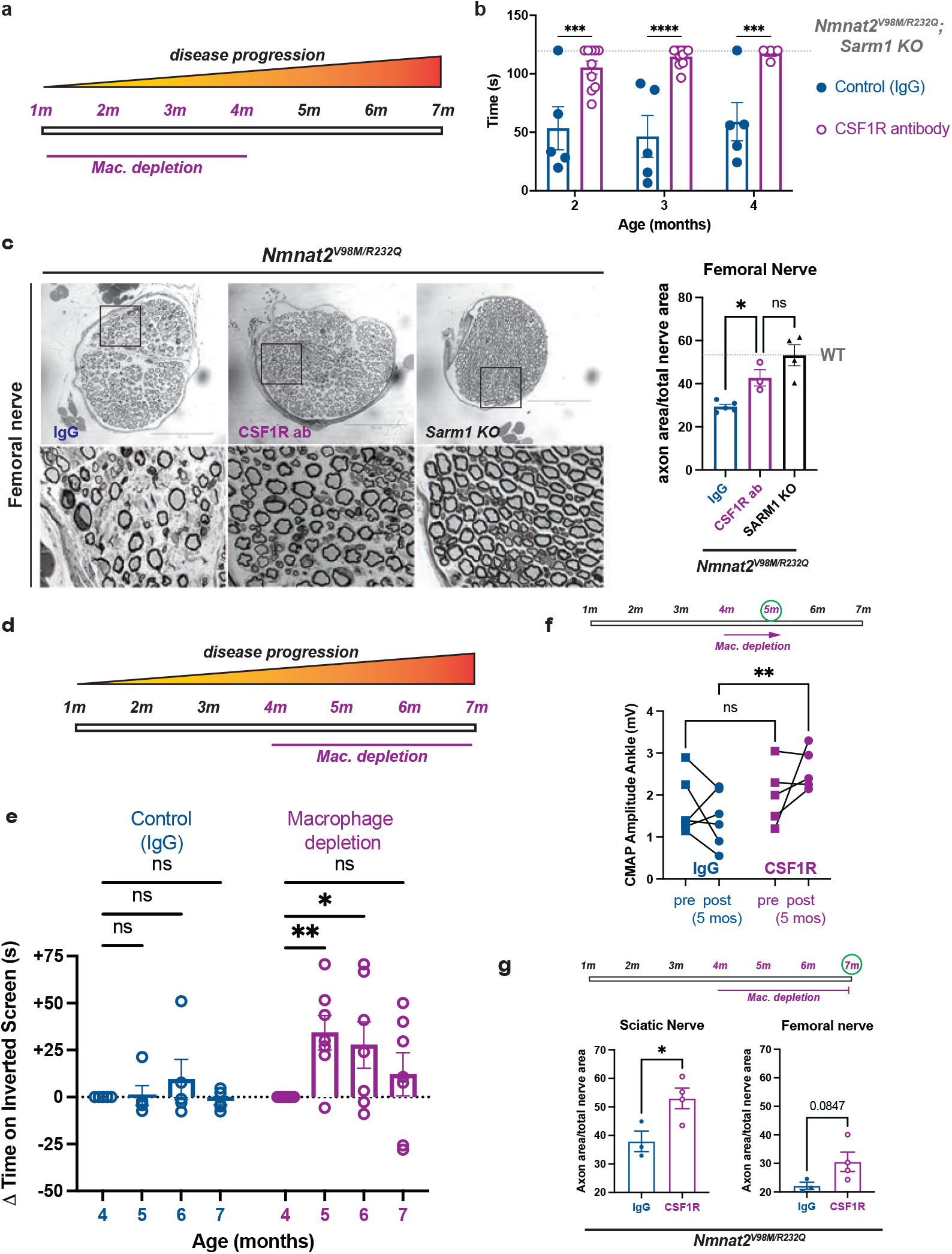
Macrophage depletion rescues motor defects and axon loss in *Nmnat2*^*V98M/R232Q*^ mice. **a**, Schematic of CSF1R antibody-mediated macrophage depletion in young (1 month) *Nmnat2*^*V98M/R232Q*^ mice. **b**, Average time suspended from an inverted screen (max. 120s) for IgG control and CSF1R-treated *Nmnat2*^*V98M/R232Q*^ mice. Statistical significance was determined by two-way ANOVA with multiple comparisons. **c**, Representative images of femoral nerve from IgG (control), CSF1R, or *Sarm1 KO Nmnat2*^*V98M/R232Q*^ mice at 4 months. Scale bar represents 150μm. Percent axonal area as a ratio of total nerve area for femoral nerve calculated to the right. **d**, Schematic of CSF1R antibody-mediated macrophage depletion in aged (4 months) *Nmnat2*^*V98M/R232Q*^ mice. **e**, Change in inverted screen time (s) from pre-treatment (4 months) measured at 5, 6, and 7 months of age comparing CSF1R treatment or IgG (control). Statistical significance was determined by two-way ANOVA with multiple comparisons. **f**, CMAP amplitude of *Nmnat2*^*V98M/R232Q*^ mice at the ankle before and after 1 month of CSF1R treatment or IgG treatment. **g**, Percent axonal area as a ratio of total nerve area for femoral nerve and sciatic nerve calculated at 7 months of age (3 months of macrophage depletion or treatment with isotype control IgG). **f-g**, Statistical significance was determined by Student’s unpaired t-test. All data are presented as mean +/- SEM. ns: not significant, *p<0.05, **p<0.01, ***p<0.001, ****p<0.0001.

Given our finding that macrophages contribute to axonal dysfunction early in the disease, we next tested whether macrophage depletion would be therapeutic after the initiation of symptoms. By four months of age, *Nmnat2*^*V98M/R232Q*^ mice exhibit profound pathological and functional motor disease (**Fig. 6b-c**). Thus, we administered the CSF1R monoclonal antibody (or control IgG) to four-month-old *Nmnat2*^*V98M/R232Q*^ mice to test whether macrophage depletion could halt or reverse disease progression (**Fig. 6d**). One month after antibody treatment, the previously symptomatic mice demonstrate a significant increase in inverted screen performance, demonstrating a profound recovery of muscle strength (**Fig. 6e**). Improvement in overall strength is accompanied by improved distal CMAP responses (**Fig. 6f**). Such a treatment response indicates that endogenous reparative processes occur after macrophage depletion to promote functional recovery. Indeed, in nerve and NMJ pathological studies, we detected ongoing axon degeneration and regeneration processes in *Nmnat2*^*V98M/R232Q*^ mice at steady-state. Rescue of muscle strength persists until the mice are 7 months old, at which point muscle strength is maintained at pre-treatment levels. Endpoint examination of macrophage-depleted peripheral nerves reveals significant rescue of sciatic nerve axon loss and a trend toward rescue of femoral nerve axon loss (**Fig. 6g**). Altogether, these data support a model wherein macrophage depletion blocks axon degeneration, tipping the balance toward innate axon reparative processes and, thus, symptom reversal. Importantly, the surprising finding that macrophage depletion can reverse both behavioral and electrophysiologic defects underscores the potential of acute macrophage-targeted therapies in chronic neurologic diseases.

## DISCUSSION

Genetic deletion of NMNAT2 is perinatal lethal; however, when combined with SARM1 deletion, mice are viable and resistant to injury-induced axon degeneration^12^. Recent advances in our understanding of the mechanisms underlying axon degeneration directly connected NMNAT2 loss to SARM1 activation through dynamic changes in the NMN-to-NAD^+^ ratio. These studies suggest that defects in NMNAT2 could predispose an individual to develop neurodegenerative disease; however, there is little direct evidence connecting mutations in the genes involved in axon degeneration to human neurological disease. Here, we examine two brothers with severe neuropathy associated with *NMNAT2* mutations. This neuropathy initially presented with sensory and motor symptoms and progressed to be a predominantly motor neuropathy with severe muscle wasting. To study the molecular mechanism underlying this syndrome, we developed a mouse model harboring the *Nmnat2*^*V98M/R232Q*^ mutations. This mutant mouse recapitulates the cardinal motor features of the human syndrome. We find that NMNAT2 enzymatic deficiency leads to chronic SARM1 activation that, in turn, leads to non-cell-autonomous macrophage activation and axon loss. Our study has several important implications. First, the creation of our mouse model enables longitudinal examination of a SARM1-dependent neuropathy, or SARMopathy, and provides a powerful platform for testing novel axo-protective therapeutics in disorders of chronic SARM1 activation. Second, we find that a major pro-degenerative role of SARM1 involves the activation of macrophages in parallel with its canonical cell-autonomous destructive functions. Finally, the identification of macrophages as key drivers of neuropathology suggests that macrophage depletion therapy could be efficacious in other diseases that involve SARM1-mediated axon degeneration.

While axon degeneration is classically thought of as a programmed injury response to acutely damaged axons, there is growing evidence that this program is aberrantly activated in progressive neurodegenerative disease. Despite the abundance of acute injury models involving SARM1 activation, a chronic model that is more akin to human progressive neurodegenerative disorders is unavailable. Such a model is necessary for testing therapeutics under conditions of subacute SARM1 activation, which our work now shows involves previously unappreciated complex biological mechanisms. We find that SARM1 is activated at an early age in *Nmnat2*^*V98M/R232Q*^ mice and that the disease is indeed fully SARM1-dependent. Moreover, this suggests that SARM1 activation and thus axon loss does not occur as an “all-or-nothing” event in chronic neurodegenerative disorders. Rather, this disorder is characterized by persistent SARM1 activity, suggesting that therapeutic intervention could be efficacious across a wide disease timeline in such disorders.

Patients with compound heterozygous variants *NMNAT2*^*V98M*^ and *NMNAT2*^*R232Q*^ develop a unique sensorimotor neuropathy involving both chronic and episodic symptomology. Chronic features of the disease are motor predominant, while episodic features involve prominent sensory and motor components. The patients typically develop acute episodes after infection, whereas the *Nmnat2*^*V98M/R232Q*^ mouse model unsurprisingly shows no evidence of episodic disease features, likely due to housing in a pathogen-free environment. The mouse model does recapitulate cardinal features of the chronic symptomatology, including a motor axonal neuropathy, distal muscle wasting and weakness, and a progressive disease course. Unlike the patients, the *Nmnat2*^*V98M/R232Q*^ mouse model has no evidence of a neuropathic pain phenotype, suggesting either species-dependent differences or that the episodic attacks seen in the patients but not in the mice contribute to the neuropathic symptoms. Interestingly, two sisters homozygous for a different variant in *NMNAT2* (T94M) that largely retains enzymatic activity also display peripheral neuropathy, albeit significantly milder than the patients described in this study^27^.

From these studies in patients with the *NMNAT2*^*V98M/R232Q*^ mutations as well as these mutant mice, it appears that the peripheral nervous system is preferentially affected by NMNAT2 dysfunction, likely due to the longer length of peripheral axons. NMNAT2 has a very short half-life and is transported from the soma to the axon, thus distal regions are likely to be more sensitive to changes in NMNAT2 activity. Hence, the peripheral defects we observe are likely a result of axon length-dependent SARM1 activation. The NMNAT2/SARM1 axon degeneration pathway functions in both sensory and motor neurons; yet curiously, *Nmnat2*^*V98M/R232Q*^ mice develop primarily a motor peripheral neuropathy. This is consistent with an *Nmnat2* hypomorphic allele that also displays predominantly peripheral motor defects^73^. In contrast, in the *Nmnat2* null mice, SARM1 drives loss of *both* sensory and motor nerves^74^, begging the question as to why motor neurons are preferentially affected in *Nmnat2*^*V98M/R232Q*^ mice? Potentially, motor neurons are more vulnerable to chronic, low-level SARM1 activation, i.e. either due to a decreased tolerance for NAD^+^ loss or increased susceptibility to macrophage-induced degeneration. In favor of this hypothesis, germline constitutive active SARM1 variants are enriched in human ALS patients^29,30^, suggesting that SARM1 primarily results in motor loss *in vivo*. Additional study of differential motor versus sensory axon susceptibility is required to answer these fundamental questions.

Studies of many common chronic neurodegenerative disorders have implicated the immune system as a key driver of disease, and activated macrophages are major contributors to tissue damage^75^. Similar to CD68^+^ microglia observed in Alzheimer’s disease patients^76^ and LPS-induced central neuroinflammation^77^, CD68^+^ macrophages are abundant in the peripheral nervous system of *Nmnat2*^*V98M/R232Q*^ mice, indicating they are activated. Importantly, macrophage activation begins early in the disease before significant axon loss has occurred, and in response to unknown signals within the neural microenvironment. We find that treating *Nmnat2*^*V98M/R232Q*^ mice with AAV expressing a dominant-negative SARM1 molecule to prevent SARM1 activation specifically in neurons prevents motor dysfunction. Altogether, these data argue that macrophage activation occurs via an extrinsic SARM1-dependent signal rather than a macrophage-autonomous program in this model.

An unexpected finding of our study is that immunodepletion of macrophages both prevents and reverses *Nmnat2-*associated motor defects. Indeed, macrophage depletion in young mice blocks the development of a motor neuropathy in *Nmnat2*^*V98M/R232Q*^ mice. Moreover, recovery of overall strength and improved nerve electrophysiology in older, macrophage-depleted *Nmnat2*^*V98M/R232Q*^ mice demonstrate that axon dysfunction can be reversed in the presence of persistent SARM1 activation; however, this effect lessens as the disease progresses. While all recent mechanistic progress on SARM1 has defined it as the central driver of a *cell-autonomous* degenerative program via its NAD^+^ hydrolase activity, our data support a new paradigm wherein chronic axonal SARM1 activation can also orchestrate non-cell-autonomous axon degeneration. Importantly, these findings indicate that macrophages are downstream effectors of SARM1 activation *in vivo*, placing SARM1 at the nexus between neuroinflammation and neurodegeneration.

So, how do macrophages induce axonal dysfunction in this syndrome? Evidence from studies of disease-associated microglia has established phagocytosis of live neurons, termed “phagoptosis”, as contributing to CNS inflammation and neurodegeneration. In the present study, the absence of motor defects or axon loss after three months of macrophage depletion in young *Nmnat2*^*V98M/R232Q*^ mice suggests that macrophages target axons that are either functional at baseline or have the ability to recover (i.e. stressed-but-viable) in the absence of macrophage attack. Pathologic evidence in the nerves and NMJs of *Nmnat2*^V98M/R232Q^ mice point to a continual process of axon degeneration and regeneration, as has been observed in other chronic neurodegenerative diseases^78,79^. Moreover, reversal of motor deficits with macrophage depletion after disease onset suggests that there is effective and ongoing regeneration once axon degeneration is blocked.

Our data support a model wherein *early* axon loss and disease symptoms are due to macrophage activation and axon engulfment. We hypothesize that macrophages are key initiators of disease in this model of a SARM1-dependent motor neuropathy, akin to their previously described roles in other common neurodegenerative diseases including MS, AD, PD, and ALS^80^. We predict that SARM1 activation eventually leads to neuron-autonomous axon degeneration that is not prevented by immunomodulatory therapy. Indeed, this may explain the waning rescue offered by macrophage depletion in older *Nmnat2*^*V98M/R232Q*^ mice. The findings herein demonstrate an acute benefit for use of currently available macrophage-targeted therapy. Given that *Sarm1* knockout and SARM1 dominant negative gene therapy both confer robust protection in this model, use of future small molecule and gene therapy SARM1 inhibitors in combination with immune targeted therapies could be optimal for long-term therapy of chronic SARMopathies.

While this study focuses on a very rare genetic cause of neuropathy, SARM1 has been implicated in an expanding number of other, more common neurodegenerative diseases including traumatic brain injury^15,18,81^, diabetic neuropathy^16^, chemotherapy-induced neuropathy^14,16,52^, glaucoma^25^, and retinal degeneration^26,82^. Importantly, we and others have identified rare hypermorphic human *SARM1* alleles in patients with ALS^29,30^. Our work suggests that in addition to SARM1 inhibition, these patients are also candidates for macrophage targeted therapies. In summary, our study implies the existence of a SARM1-dependent, non-cell-autonomous pathway for axonal dysfunction that is amenable to targeted immunomodulatory therapy and presents a mechanistically defined mouse model of a pure SARMopathy in which to test such treatment strategies.

## MATERIALS AND METHODS

### Whole-exome sequencing

Whole exome sequencing and sequence analysis were performed by the Division of Genomic Diagnostics Genetic Counseling Core at Children’s Hospital of Philadelphia. Genomic DNA was extracted from peripheral blood following standard DNA extraction protocols. Targeted exons were captured with the Agilent SureSelect XT Clinical Research Exome kit per manufacturer’s protocol, sequenced on the Illumina HiSeq 2000 or 2500 platform with 100bp paired-end reads, and sequencing variants were identified using an in-house custom-built bioinformatics pipeline as described previously^83^. Mapping and analysis were based on the human genome build UCSC hg19 reference sequence. Single nucleotide variants, small deletions, and small insertions were detected. Suspected pathogenic variants were confirmed by Sanger sequencing.

### Constructs

The Takara HD InFusion Cloning Kit was used to introduce R232Q and V98M mutations into the complete open reading frame of the canonical 307 amino acid human *NMNAT2* reference allele fused to a FLAG tag followed by a 10 amino acid linker sequence and C-terminal 2 X Strep Tag cloned into lentivirus vector FCIV.

### HEK293T transfection

HEK293T cells were cultured in DMEM with 2 mM glutamine and 1% penicillin/streptomycin (both Gibco), and 10% fetal bovine serum. Cells were plated in 12-well format to reach 25-35% confluence before transfection with polyethyleneimine (PEI, 1 mg/mL, pH 7.0) using a ratio of 3:1 (mg of PEI vs. mg of plasmid DNA). 200 ng StrepTag-NMNAT2-Flag expression vectors (reference allele, V98M, or R232Q) were transfected per well. 1 mg/mL Cycloheximide (Sigma-Aldrich) was used to block protein synthesis 24 hours after transfection. Cells from a single well at each time point after treatment were suspended in 100 mL of 100 mM Tris-Cl, pH 8.0, and 150 mM NaCl with Protease Inhibitor Cocktail (Pierce, Buffer 1), sonicated to fragment genomic DNA, then mixed with 30 mL of 4x NuPage LDS sample buffer (Invitrogen). After heating to 90°C for 5 minutes, equal amounts were used for Western blot.

### Western Blot

Cell lysates were resolved using SDS polyacrylamide gel electrophoresis (PAGE) on 4-12% Bis-Tris Plus gel (Invitrogen), transferred to PVDF membrane (Invitrogen), and followed by immunoblotting for Flag (1:1000 Mouse Anti-Flag M2 monoclonal, Sigma F3165) and beta-tubulin (1:1000 Anti-beta-tubulin DSHB Clone E7), and visualization by standard chemiluminescence.

### NMNAT recombinant protein expression and purification

Constructs described above, human reference allele of *NMNAT2* and variants V98M or R232Q, were transfected to 150 mm diameter cell culture dish with ∼50% confluence of HEK293T cells. 15 mg of plasmid was mixed with 45 mg of PEI and transfected into the cells. Cells were harvested 48 hours after transfection and resuspended in Buffer 1 before lysis by sonication on ice. After centrifugation at 18,000xg for 10 minutes 3 times to remove the cell debris, the supernatant was mixed with PureCube HiCap Streptactin MagBeads (Cube Biotech) for 1 hour. Proteins were eluted with Buffer 1 plus 5 mM desthiobiotin and stored at -80 °C. Their concentrations were determined by the Bio-Rad protein assay. Protein purity was assessed by 4-12% SDS-PAGE and Coomassie staining.

### NMNAT Activity Assay

NMNAT activity was defined as 1 mmol NAD^+^ generated per min (one unit, U) per mg of protein. Typically, the reaction was initiated by mixing purified NMNAT2 protein at 1-2 mg/mL with 100 mM ATP and 100 mM NMN in 10 mM HEPES, pH 7.4, and 5 mM MgCl2 at 37 °C. At various time points, reactions were stopped by removing 100 µL from the reaction and mixing it with 100 µL 0.5 M perchloric acid (HClO4) on ice for 10 minutes. After centrifugation at 18,000xg for 10 minutes, the supernatant was mixed with 11 µL 3 M K2CO3 for neutralization. Samples were placed on ice for another 10 minutes and centrifuged. 45 µL of supernatant containing extracted metabolites was mixed with 5 µL 0.5 M Potassium Phosphate buffer and quantified by HPLC (Nexera X2) on a Kinetex (100 × 3 mm, 2.6 µm; Phenomenex) column. An NAD^+^ standard (Sigma) was used to generate a standard curve for the quantification of NAD^+^ in the extraction. NAD^+^ production at 10 minutes, which is within the steady-state, was used to calculate NMNAT activity.

### Enzymatic and half-life assay statistics

Data are presented as Mean ± SEM. Fitting of data was performed using Excel (Microsoft) or Prism 9 (GraphPad). Curve fittings and specific tests used are described in the Figure legends. A *p-value* <0.05 was considered significant.

### Generation of *Nmnat2*^*V98M/R232Q*^ compound heterozygous mice

All animal experiments were performed under the direction of institutional animal study guidelines at the Washington University, St. Louis, MO. To generate *Nmnat2*^*V98M/R232Q*^ mice via CRISPR/Cas9, guide RNAs were designed by the Washington University School of Medicine Genome Engineering & iPSC Center, and individually created gRNAs were microinjected into B6CBAF2/J pronuclei along with Cas9 protein and donor DNA oligonucleotides engineered to introduce the desired mutations by homology-directed repair. Properly mutated transgenic mice with individual mutations were confirmed by sequencing. Founder mice were initially mated to C57BL/6J mice and heterozygotes for each allele were subsequently mated together to generate trans-heterozygotes.

### Nerve structural analysis

Sciatic, sural, and femoral nerves were processed as previously described^14,84^. Briefly, nerves were fixed in 3% glutaraldehyde in 0.1 ml PBS (Polysciences) overnight at 4°C, washed, then stained with 1% osmium tetroxide (Sigma Aldrich) overnight at 4°C. Nerves were then washed, and dehydrated in a serial gradient of ethanol from 50 to 100%. After dehydration, nerves were incubated in 50% propylene oxide/50% ethanol, then 100% propylene oxide. Subsequently, nerves were incubated in the Araldite resin solution/propylene oxide solutions overnight in the following ratios: 50:50, 70:30, 90:10, and embedded in 100% Araldite resin solution (Araldite: DDSA: DMP30; 12:9:1; Electron Microscopy Sciences) and baked overnight at 60°C. Sciatic nerves were embedded so that the most distal portion was sectioned. For the light microscope analysis of 400– 600 nm semithin sections were cut using Leica EM UC7 Ultramicrotome and placed onto microscopy slides. Toluidine blue staining and quantification were performed as previously described^84^. All quantifications were performed by an individual blinded to genotype.

### Tissue immunohistochemistry

#### Nervous tissue

Six micron-thick sections of sciatic nerves and twenty micron-thick sections of spinal cord were prepared on a cryostat (Leica CM1860), mounted onto slides, and processed as described^14^. For visualization of activated macrophages, the primary antibody used was 1:100 CD68 (rat, Bio-Rad), and the secondary antibody was anti-rat Cy3 (Jackson Immunoresearch) at a dilution of 1:500. For visualization and quantification of GFP fluorescence in spinal cords, directly conjugated rabbit anti-GFP Alexa 488 (Invitrogen) was used 1:200. For visualization of motor neuron cell bodies, the primary antibody used was 1:100 ChAT (rat, Millipore), and the secondary antibody was 1:250 anti-goat-Cy3 (Jackson Immunoresearch). Sections were cover-slipped with Vectashield with DAPI (Vector Laboratories) to allow visualization of nuclei. Sciatic nerves were imaged with the Leica DMI 4000B confocal microscope using a 40× immersion oil. Spinal cords were imaged at 10x magnification using a scheme covering 100% of the total spinal cord area with image stitching for quantification. Z-stacks were acquired through the whole sample, max projected, then exported for quantification. ImageJ was used for quantification of integrated fluorescence density quantification from three independent sections per mouse. Motor neuron numbers were manually counted in the ventral horn using ImageJ (3-4 mice per genotype).

#### Tibialis Anterior Muscle

Ten micron-thick sections of the tibialis anterior muscle were processed as described^14^. The primary laminin antibody was used at 1:100 (Sigma #L9393). The secondary antibody was 1:500 anti-rabbit Cy3 (Jackson Immunoresearch). Muscle fiber cross-sectional area was calculated by averaging fiber cross-sectional area from ≥150 muscle fibers per mouse from three independent 20X images. Three mice were used per genotype in each age cohort.

#### Lumbricals

The mice were transcardially perfused with 20ml of 4% paraformaldehyde (PFA) solution. Dissected lumbrical muscles were permeabilized with 2% TritonX-100 /1x PBS (PBST) for 30min and then blocked with 4% bovine serum albumin (BSA) dissolved in 0.3% PBST 30mins at room temperature. Muscles were incubated with primary antibodies of SV2 (1:200, DSHB AB2315387) and 2H3 (1:100, DSHB AB2314897) overnight at 4°C. After incubation of primary antibody, muscles were incubated with FITC rabbit anti-mouse IgG1 (1:400, Invitrogen A21121) and Alexa fluoro-568 conjugated α-bungarotoxin (1;500, biotium 00006) for 2hrs at room temperature. Muscles were washed with PBS for 5 min 3 times and then were mounted by mounting media (VECTASHIELD).

To analyze NMJ morphology, z-stack images were obtained. Maximal intensity projection images were analyzed to determine the volume of endplates using Imaris software. 15-20 NMJs were analyzed per mouse. NMJ occupancy was calculated as a ratio of pre-synaptic axonal area (SV2/2H3) versus post-synaptic area (BTX). As the value approaches 1.0, it indicates motor endplates are fully occupied by axons.

### Intraepidermal nerve fiber density quantification

Intraepidermal nerve fiber (IENF) staining and quantification were performed as previously described^14,85^. Briefly, footpad skin was dissected and then placed in freshly prepared Zamboni’s fixative overnight at 4°C. Samples were then thoroughly washed with PBS then incubated for 24 hr in 30% sucrose in PBS. Samples were embedded in O.C.T. then sectioned at 50 μm using a cryostat. Sections were stored −20°C in a cryoprotectant comprised of 30% sucrose and 33% ethylene glycol in PBS until ready for immunohistochemistry. Free floating sections were washed with PBS, blocked with 5% normal goat serum in PBST, then incubated in PGP9.5 (AB1761, Millipore) diluted 1:1000 in blocking buffer overnight at 4°C. Free floating sections were washed with PBST, incubated with species appropriate secondary antibody (anti-rabbit-Cy3) 1:500 for 2 hr at room temperature, washed with PBST, and then mounted in Vectashield with DAPI.

The footpads were identified and imaged on a Leica DMI 4000B confocal microscope using a 10× objective. Z-stacks were acquired through the whole sample, max projected, then exported for quantification. IENF density was quantified as the number of PGP9.5^+^ axons that crossed the basement membrane normalized to the length of the basement membrane. IENF densities were averaged in three separate sections for each animal. Imaging and analysis were performed by an individual blinded to genotype.

### Tissue metabolite measurements

On the day of metabolite extraction, sciatic nerve tissues were homogenized in 160 µL of cold 50% MeOH solution in water using a homogenizer (Branson) and then centrifuged (15,000 g, 4°C, 10 min). The clear supernatant was transferred to a new tube containing 50 µL chloroform and vigorously shaken then centrifuged (15,000 g, 4°C, 10 min). The chloroform extraction was repeated twice. The clear aqueous phase (100µL) was transferred to the new tube and then lyophilized and stored at 80°C until measurement. Lyophilized samples were reconstituted with 50 µL of 5 mM ammonium formate (Sigma) and centrifuged at 12,000 x g for 10 min. Cleared supernatant was transferred to the sample tray. Nervous tissue metabolite measurements were acquired as previously described^84^.

### AAV constructs and virus injections

AAV constructs were created as previously described^22^. Briefly, AAV8-hSYN-SARM1-DN-EGFP was generated by the viral vector core of the Hope Center for Neurological Disorders at Washington University in St. Louis. Viral particles were purified by iodixanol gradient ultracentrifugation, and virus titers were measured by dot blot. Under light anesthesia with Avertin, 6 × 10^11^ viral genomes were injected intrathecally at L6/S1. Viral expression was confirmed by detecting EGFP expression via immunohistochemical analysis of sectioned spinal cords.

### Nerve electrophysiology

Compound muscle action potentials (CMAPs) were acquired as previously described^86^ using a Viking Quest electromyography device (Nicolet). Mice were anesthetized, then electrodes (stimulating: ankle or sciatic notch; recording: footpad) were put into place. Supramaximal stimulation was used for CMAPs. SNAPs were acquired as previously described^14^ using a Viking Quest electromyography device (Nicolet). Briefly, mice were anesthetized, then electrodes were inserted subcutaneously into the tail (recording electrode at the base, stimulating electrodes 30 mm distal from the negative recording electrode, and ground between the stimulating and recording electrodes). Supramaximal stimulation was used for SNAPs.

### Behavioral tests

#### Tail flick assay

The tail flick assay was performed as previously described^87^. Briefly, mice were restrained horizontally, and the tip of their tails was submerged in a 55°C water bath. All mice removed their tails from the noxious stimulus by flicking their tails out of the water. The latency to withdrawal from the water bath was measured using a stopwatch. Five trials were performed and averaged for each mouse.

#### Inverted screen assay

The inverted screen assay was performed as previously described with minor modifications^88^. Mice were placed on a wire mesh screen. The latency to fall for each mouse was recorded, and each mouse underwent three trials with five-minute rest periods. If a mouse did not fall off the mesh screen within 120 seconds, then that time was recorded, and the mouse was taken off the screen.

#### Hindlimb grip strength

Hindlimb grip strength was measured using a computerized grip strength meter. To measure hindlimb grip strength, mice were held upright at the base of the neck and the tail, and the mouse was held over a bar connected to a force transducer. Once the mouse gripped the bar with its hind paws, the mouse was pulled backward until the grip was released. The maximum force of each measurement was measured in grams and recorded by the grip strength meter. Each mouse underwent five trials. If a mouse could not establish a grip on the bar, then the value for that trial was recorded as zero.

### Flow cytometry

Nerves were collected and kept on ice until dissociation. Cells were then minced and incubated with gentle shaking for 20 min in digestion media containing 0.3% collagenase IV, 0.04% hyaluronidase, and 0.04% Dnase in DMEM at 37 °C. Cells were then washed and filtered through 70μm cell strainers. Single-cell suspensions were stained at 4 °C. Dead cells were excluded by propidium iodide (PI). Antibodies to the following proteins were used: CD11b (clone M1/70), CD45 (clone 30-F11), and CD64 (clone X54-5/7.1). Cells were analyzed on an LSRII flow cytometer (Becton Dickinson) and analyzed with FlowJo software.

### RNA Sequencing and Analysis

Samples were prepared according to library kit manufacturer’s protocol, indexed, pooled, and sequenced on an Illumina NovaSeq 6000. Basecalls and demultiplexing were performed with Illumina’s bcl2fastq software and a custom python demultiplexing program with a maximum of one mismatch in the indexing read. RNA-seq reads were then aligned to the Ensembl release 76 primary assembly with STAR version 2.5.1a^1^. Gene counts were derived from the number of uniquely aligned unambiguous reads by Subread:featureCount version 1.4.6-p5^2^. Isoform expression of known Ensembl transcripts were estimated with Salmon version 0.8.2^3^. Sequencing performance was assessed for the total number of aligned reads, total number of uniquely aligned reads, and features detected. The ribosomal fraction, known junction saturation, and read distribution over known gene models were quantified with RSeQC version 2.6.2^4^.

All gene counts were then imported into the R/Bioconductor package EdgeR^5^ and TMM normalization size factors were calculated to adjust for samples for differences in library size. Ribosomal genes and genes not expressed in the smallest group size minus one samples greater than one count-per-million were excluded from further analysis. The TMM size factors and the matrix of counts were then imported into the R/Bioconductor package Limma^6^. Weighted likelihoods based on the observed mean-variance relationship of every gene and sample were then calculated for all samples and the count matrix was transformed to moderated log 2 counts-per-million with Limma’s voomWithQualityWeights^7^. The performance of all genes was assessed with plots of the residual standard deviation of every gene to their average log-count with a robustly fitted trend line of the residuals. Differential expression analysis was then performed to analyze for differences between conditions and the results were filtered for only those genes with Benjamini-Hochberg false-discovery rate adjusted p-values less than or equal to 0.05.

For each contrast extracted with Limma, global perturbations in known Gene Ontology (GO) terms, MSigDb, and KEGG pathways were detected using the R/Bioconductor package GAGE^8^ to test for changes in expression of the reported log 2 fold-changes reported by Limma in each term versus the background log 2 fold-changes of all genes found outside the respective term. The R/Bioconductor package heatmap3^9^ was used to display heatmaps across groups of samples for each GO or MSigDb term with a Benjamini-Hochberg false-discovery rate adjusted p-value less than or equal to 0.05. Perturbed KEGG pathways where the observed log 2 fold-changes of genes within the term were significantly perturbed in a single-direction versus background or in any direction compared to other genes within a given term with p-values less than or equal to 0.05 were rendered as annotated KEGG graphs with the R/Bioconductor package Pathview^10^.

To find the most critical genes, the Limma voomWithQualityWeights transformed log 2 counts-per-million expression data was then analyzed via weighted gene correlation network analysis with the R/Bioconductor package WGCNA^11^. Briefly, all genes were correlated across each other by Pearson correlations and clustered by expression similarity into unsigned modules using a power threshold empirically determined from the data. An eigengene was then created for each de novo cluster and its expression profile was then correlated across all coefficients of the model matrix. Because these clusters of genes were created by expression profile rather than known functional similarity, the clustered modules were given the names of random colors where grey is the only module that has any pre-existing definition of containing genes that do not cluster well with others. These de-novo clustered genes were then tested for functional enrichment of known GO terms with hypergeometric tests available in the R/Bioconductor package clusterProfiler^12^. Significant terms with Benjamini-Hochberg adjusted p-values less than 0.05 were then collapsed by similarity into clusterProfiler category network plots to display the most significant terms for each module of hub genes in order to interpolate the function of each significant module. The information for all clustered genes for each module was then combined with their respective statistical significance results from Limma to determine whether or not those features were also found to be significantly differentially expressed.

### CSF1R antibody injections

Macrophages were depleted by administering 1.5 mg of anti-mouse CSF1r (clone AFS98, BioXCell) or Rat IgG (cat #I4131, Sigma) i.p. every three weeks for a total of four injections. Depletion of macrophages was confirmed by CD68 immunofluorescence in the sciatic nerve.

### Macrophage activation assay

Peritoneal macrophages were harvested from individual mice by peritoneal lavage using ice-cold DMEM containing 1% FBS. Cells were then transferred into plates and subjected to either PBS or LPS treatment at a concentration of 10 ng/mL for 4 hours at 37 °C.

### Statistical analysis

Unless otherwise stated, data are reported as means ± standard error of the mean (SEM). Between group comparisons were made with one-way and two-way ANOVA with post hoc Holm-Sidak multiple comparison test or paired and unpaired t-tests as appropriate. Two-sided significance tests were used throughout and P<0.05 was considered statistically significant. All statistics were calculated with the aid of Prism 9 (GraphPad) software.

## Acknowledgments

We thank the patients and their family for participating in our research study. We thank Milbrandt and DiAntonio labs members for helpful discussions and Cassidy Menendez, Rachel McClarney, Timothy Fahrner, and Alicia Neiner for technical support. We also thank the CHOP Genetic Counseling Core and the Genome Engineering and iPSC Center (GEIC).

## Author Contributions

C.B.D., A.J.B., A.D., and J.M. conceived the overall study. All authors contributed to the study design. C.B.D. and A.S. collected and analyzed all mouse behavior data and mouse tissue samples. J.Z. performed *in vitro* biochemical experiments. S.W.Y. provided and interpreted clinical and genetic information. R.E.S. performed ultrastructural electron microscopy nerve analysis. P.L.W. and C.B.D. performed and analyzed flow cytometry experiments. Y.Y. performed NMJ analysis. A.Y.K. analyzed bulk RNA-Seq data. Y.S. provided early intellectual input. C.B.D. wrote the manuscript and prepared all figures. A.J.B., A.D., and J.M. oversaw the analysis and revised the manuscript. All authors gave final approval of the manuscript.

## Corresponding authors

Correspondence to J.M. and A.D.

## Funding

This work was supported by National Institutes of Health grants (R01NS119812 to A.J.B., A.D. and J.M, R01NS087632 to A.D. and J.M., R37NS065053 to A.D., and RF1AG013730 to J.M.)

## Competing Interests

A.D. and J.M. are co-founders, scientific advisory board members, and shareholders of Disarm Therapeutics, a wholly-owned subsidiary of Eli Lilly. A.J.B. and Y.S. are consultants to Disarm Therapeutics. The authors have no other competing conflicts or financial interests.

---

**Supplemental Figure 1:**
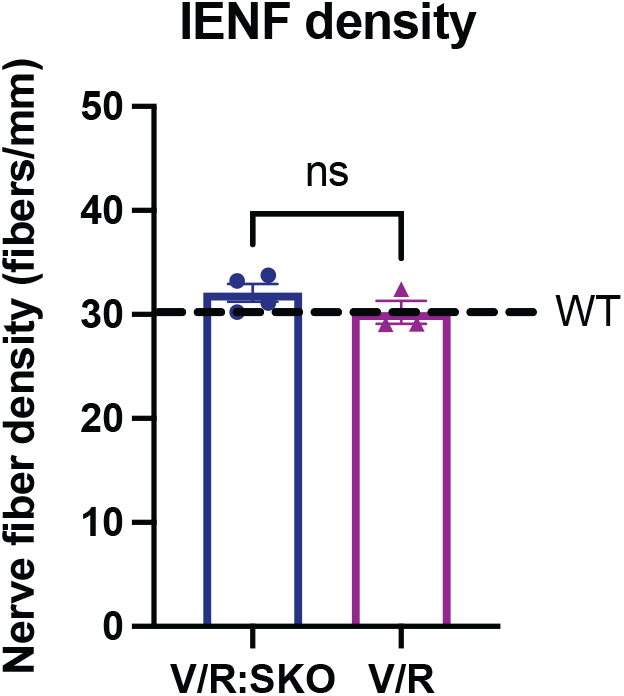
Intraepidermal nerve fiber density is unaffected in *Nmnat2*^*V98M/R232Q*^ mice. Intraepidermal nerve fiber density of footpad skin of 12-month-old *Nmnat2*^*V98M/R232Q*^; *Sarm1 KO* and *Nmnat2*^*V98M/R232Q*^ mice.

**Supplemental Figure 2:**
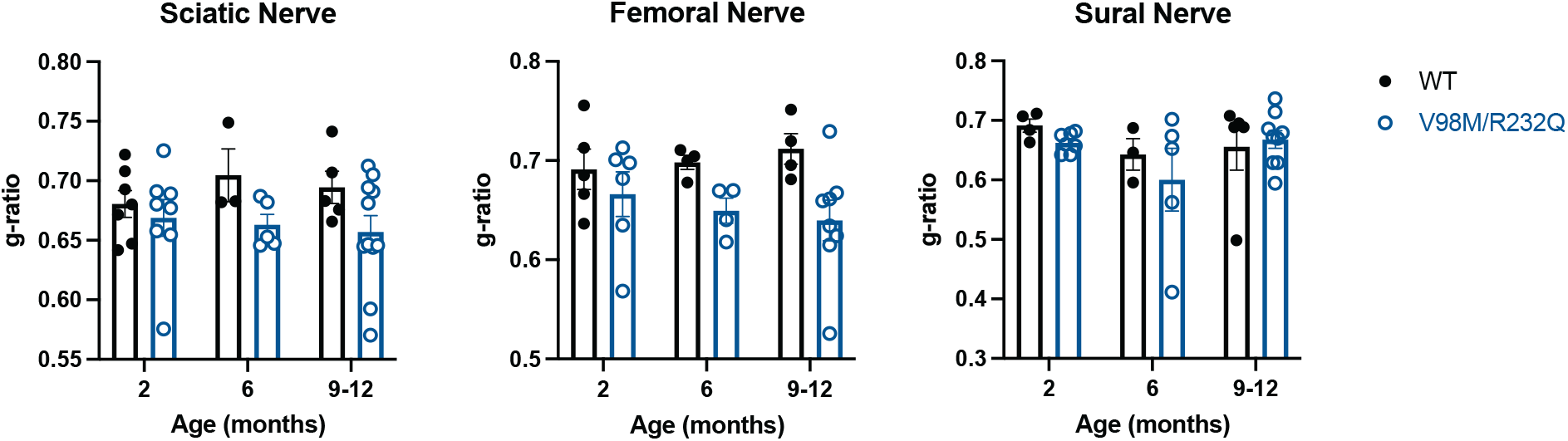
Myelin thickness is preserved in *Nmnat2*^*V98M/R232Q*^ mice. Average g-ratio in sciatic, femoral, and sural nerves at 2, 6, and 9-12 months of age, comparing *Nmnat2*^*V98M/R232Q*^ mice to WT mice. All data are presented as mean +/- SEM.

**Supplemental Figure 3:**
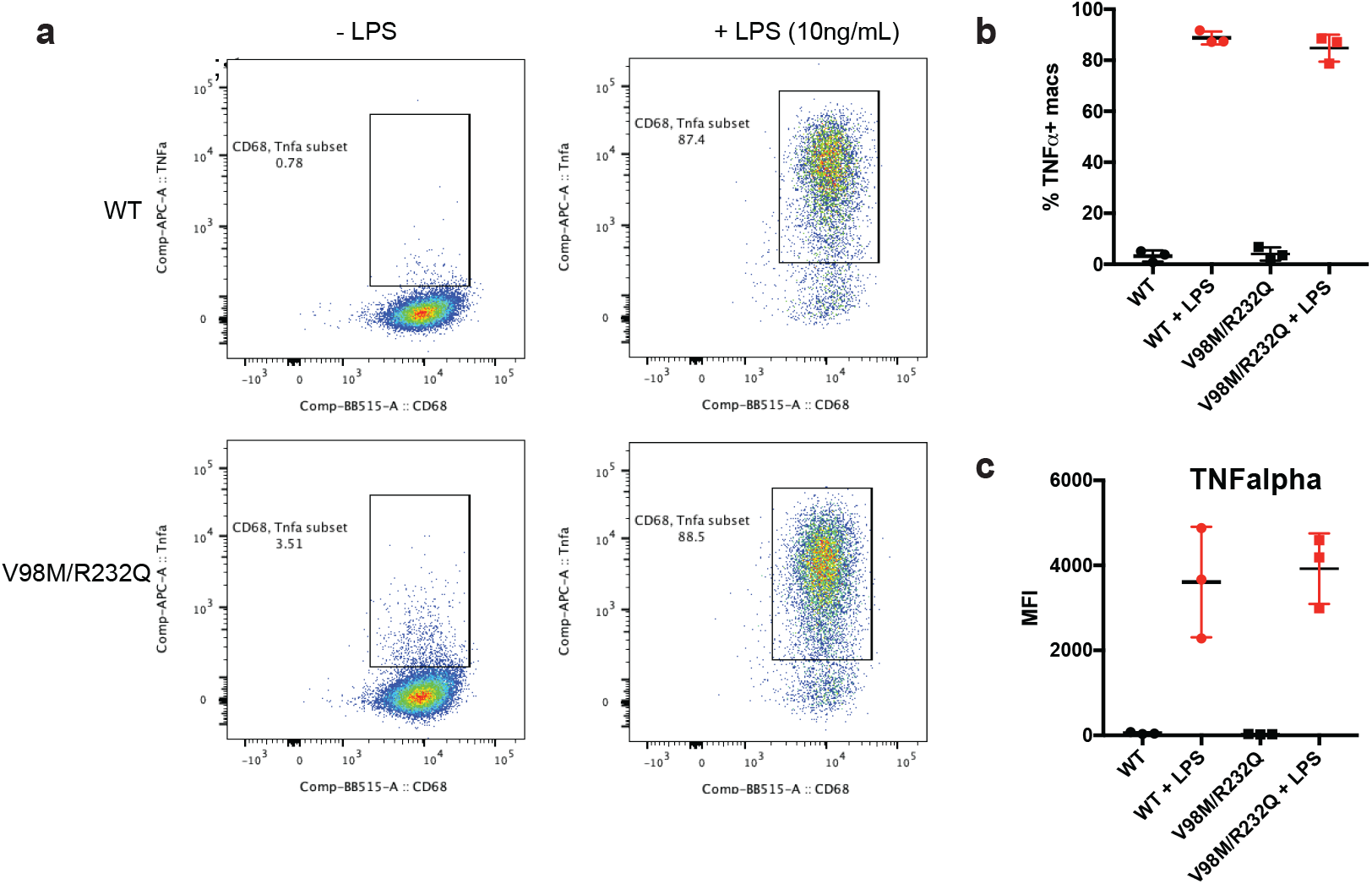
Macrophage activation assay. **a-b**, Representative scatter plots (a) and quantification (b) of CD68^+^ TNF-alpha^+^ *Nmnat2*^*V98M/R232Q*^ or WT peritoneal macrophages in the presence or absence of LPS (10ng/mL) stimulation for 4hrs. **c**, Mean fluorescence intensity (MFI) of TNF-alpha cell surface expression on CD68^+^TNFalpha^+^ peritoneal macrophages in the presence or absence of LPS (10ng/mL) stimulation for 4hrs.

**Supplemental Figure 4:**
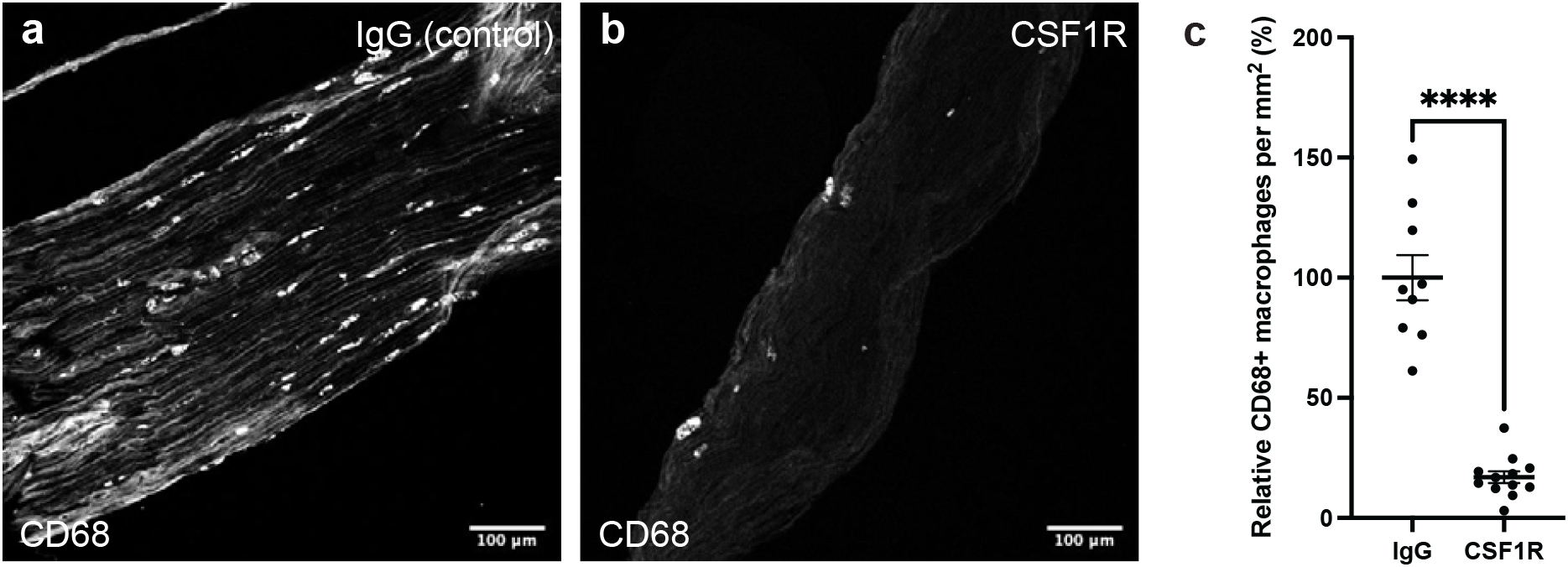
Macrophage depletion quantification. **a-b**, Representative images of CD68^+^ cells in sciatic nerves after 3 months of CSF1R (a) or IgG (b) injections. **c**, Quantification of CD68^+^ cells per mm^2^ nerve area relative to IgG control nerves (reported as a percentage of control).

